# Mechanistic pathways of tick exposure risk in native and invaded plant communities

**DOI:** 10.1101/2024.11.19.624319

**Authors:** Drew Hiatt, Whalen W. Dillon, Allison Gardner, Steven Cabrera, Michael Dietze, Brian F. Allan, S. Luke Flory

## Abstract

Plant invasions may alter disease vector abundance by several mechanistic pathways, including modifying microclimates that influence vector survival or changing habitats to influence host use. Here, we used a field experiment and observational data to evaluate multiple mechanistic pathways (tick survival and host abundance) by which plant invasions may alter vector-borne disease risk using the common disease vector lone star tick (*Amblyomma americanum*), its preeminent host white-tailed deer (*Odocoileus virginianus*), and the widespread invasive cogongrass (*Imperata cylindrica*) in the southeastern USA. In the field experiment, ticks survived over 50% longer in areas dominated by the invasive plant compared to those with only native plant species. Invaded areas had lower temperatures and higher relative humidity, yielding a lower vapor pressure deficit (VPD) that likely reduced tick desiccation. The observational study showed similar average tick abundance in native and invaded plant communities and no difference in wildlife host (white-tailed deer) activity between plant communities. However, there was a positive relationship between tick abundance and white-tailed deer activity, but only in native areas. Together, these results suggest that more favorable microclimate conditions resulting in greater tick longevity are the dominant driver of tick abundance in invaded areas, while tick abundance in native-dominated areas may be promoted, at least in part, by white-tailed deer activity. Our results demonstrate that plant invasions can affect multiple, potentially counteracting mechanistic pathways that contribute to tick exposure risk. The complexity of these relationships highlights the need for better understanding of how invasive species and other global change drivers influence disease vectors and, ultimately, disease transmission.

## Introduction

Human-mediated introductions of species to non-native ranges have resulted in widespread biological invasions, which are a key driver of global environmental change (Vitousek et al. 1996). Accumulating evidence demonstrates that invasive species can have direct effects on ecological communities by altering, for example, biodiversity, nutrient cycling, and disturbance regimes (Vilà et al. 2011, Pyšek et al. 2012). However, they can elicit indirect effects that may be larger than their direct effects, such as through changes to habitat structure and microclimate (Linders et al. 2019). These indirect effects can have cascading impacts on humans by affecting food production, recreation, and climate mitigation (Pejchar and Mooney 2009). Invasive species may also indirectly influence human health if they alter interactions among species that transmit diseases to humans (Rai and Singh 2020), however, to our knowledge such effects have been rarely explored.

Infectious pathogens transmitted from animals to humans via arthropod vectors such as mosquitoes and ticks place a major burden on global health (Morens et al. 2004, Jones et al. 2008). Tick-borne disease transmission to humans results from complex species interactions that are mediated by impacts of plant communities on tick growth, survival, dispersal, questing, and reproduction (Ostfeld et al. 2018, Morand and Lajaunie 2021) and host habitat conditions that affect the frequency and duration of host use (Elias et al. 2006, Allan et al. 2010a, Guiden and Orrock 2019). Plant invasions can alter multiple mechanistic pathways that affect tick abundance and infectious disease exposure for humans by modifying either habitats that alter host abundance or microclimate conditions that promote or inhibit tick survival (Allan et al. 2010a, Williams and Ward 2010, Stewart et al. 2021) (Figure 1). Entomological metrics of exposure risk include measures of tick abundance (e.g., density of ticks), pathogen infection prevalence, or their product (e.g., density of infected ticks) (Fischhoff et al. 2019). Vector abundance often is used as a proxy for tick exposure risk, indicating a human’s likelihood of encountering a tick upon entering a particular habitat (Barbour and Fish 1993, Fischhoff et al. 2019). While there has been research on how these pathways individually affect tick abundance, studies that simultaneously address multiple pathways are needed to improve understanding of complex processes by which invasive plants may alter tick exposure risk, and thus infectious disease.

**Figure 1.**
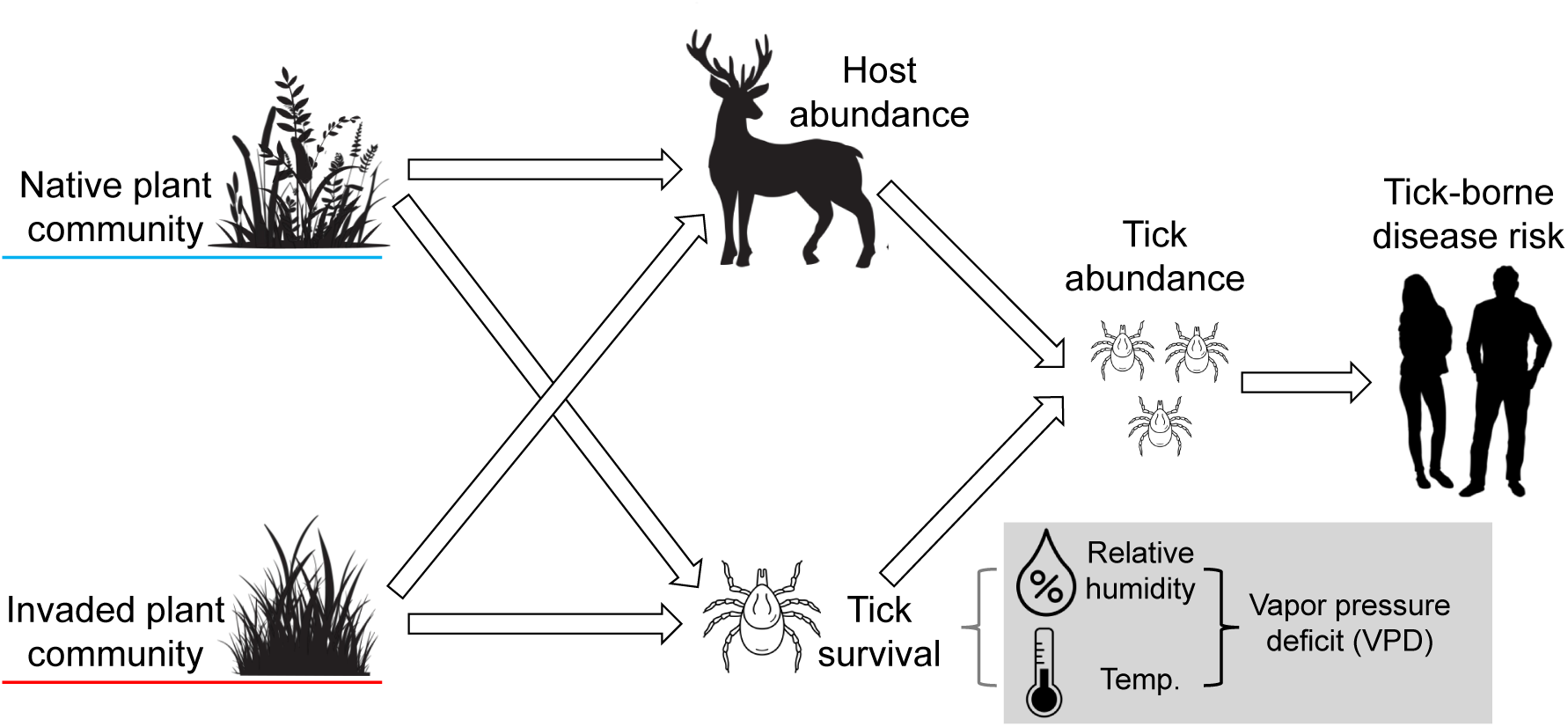
Conceptual diagram of two pathways that contribute to tick-borne disease risk. Native and invaded plant communities may have different effects on both host (e.g., deer, rodent) abundances and tick survival, which together determine tick abundance and, in turn, influence human tick exposure and tick-borne disease risk. Differences between invaded and native areas in microhabitat variables such as relative humidity and temperature, which determine vapor pressure deficit (VPD), can affect off-host tick survival. Vector graphics are copyright-free images obtained from stock.adobe.com via the University of Florida’s enterprise license.

Collectively, evidence for mechanisms that influence tick-borne disease in invaded areas is sparse and inconsistent, and varies by tick species, invasive plant species, and habitat conditions, among other factors (Civitello et al. 2008, Allan et al. 2010a, Williams and Ward 2010, Mack and Smith 2011, Racelis et al. 2012, Adalsteinsson et al. 2016). Plant invaders have been shown to increase habitat use by small mammals (Guiden and Orrock 2019), and white-tailed deer (Elias et al. 2006, Allan et al. 2010a), leading to greater abundance of ticks (Elias et al. 2006) and pathogen-infected ticks (Allan et al. 2010a) compared to native areas. Plant invasions may also increase tick abundances relative to native plant dominated areas by providing favorable microclimate conditions (i.e., higher humidity, lower temperature, or lower vapor pressure deficit) that reduce tick desiccation (Williams and Ward 2010, Racelis et al. 2012, Brunner et al. 2023). However, the few studies comparing the abiotic mechanisms affecting tick survival in invaded and native areas show either no difference (Allan et al. 2010a) or lower tick survival (Civitello et al. 2008) in invaded compared to native plant dominated communities. Given the ongoing introduction and spread of both invasive plants (Seebens et al. 2017), and ticks (Ostfeld and Brunner 2015), there is a critical need to better understand how invasive plants affect host habitat use (Stewart et al. 2021) as well as the abiotic factors that influence tick survival.

Here, we measured differences in tick abundance in native and invaded plant communities, and how these may be attributable to tick survival and host activity, which are the two main pathways that determine tick abundance (Figure 1), in the two plant community types. Specifically, we used field experiments and observational field studies to compare microclimate conditions and tick survival (lone star tick, *Amblyomma americanum*) in native plant dominated areas compared to areas invaded by cogongrass (*Imperata cylindrica*). We also measured relative activity of a key host, white-tailed deer (*Odocoileus virginianus*), in invaded and native areas, and evaluated how tick abundance was related to host activity. White-tailed deer serve as an important host for all three life stages of *A. americanum* (Bloemer et al. 1986).

First, in a field experiment, we measured survival of lab-reared ticks in native plant communities and adjacent invaded areas while concurrently measuring temperature and relative humidity (RH) to calculate vapor pressure deficit (VPD), a measure of the strength of atmospheric desiccation (Grossiord et al. 2020). Second, we evaluated relative white-tailed deer activity (via camera traps) and abundances of host-seeking ticks (via CO_2_ trapping) in native and invaded sites in a field survey. Because the invader creates much denser stands compared to native plant communities (Appendix S1: Figure S1), we hypothesized that invaded areas would have lower temperatures and higher relative humidity, yielding lower vapor pressure deficit, which would be more favorable for tick survival relative to native areas (Alba et al. 2017). We also hypothesized that relative white-tailed deer use would be greater in invaded communities because dense cover created by the invader would provide perceived cover from predators, as documented with small mammals in experimental *Imperata*-invaded plots (Borden et al. 2022), which would in turn be linked to greater tick abundance.

## Methods

### Study system

One of the most widely distributed invasive plant species in the southeastern USA is *Imperata cylindrica,* (L.) Beauv. (cogongrass; hereafter *Imperata*) (Estrada and Flory 2015), and it is one of the top five most costly terrestrial invasive plants to manage in Florida (Hiatt et al. 2019). *Imperata* can invade a wide variety of habitats due in part to its greater trait plasticity (Hiatt and Flory 2020), and drought tolerance (Fahey et al. 2018) compared to co-occurring native plant species. *Imperata* invasions can alter the structure of southern pine forests by producing greater fire intensity compared to native plant communities (Dillon et al. 2021), leading to enhanced fire induced tree mortality (Flory et al. 2022). *Imperata* has also been shown to change microclimate conditions, including greater soil moisture and relative humidity, and lower temperatures at ground level compared to native plant communities due to its higher stem density and greater leaf area (Alba et al. 2017, Cabrera et al. 2021).

Lone star ticks (*Amblyomma americanum*) are hard ticks in the family Ixodidae (Keirans and Litwak 1989) that can transmit pathogens, including the causative agents of human ehrlichiosis, Rocky Mountain spotted fever, and tularemia (Childs and Paddock 2003). Time between blood meals (off-host), either seeking hosts to parasitize, digesting blood meals, or dormant, comprises over 90% of an Ixodid tick’s life cycle, making desiccation (i.e., loss of water through the cuticle) (Needham and Teel 1991) the primary cause of off-host tick mortality (Ostfeld and Brunner 2015). Maintaining a constant water balance is one of the most important processes that influence off-host survival and, ultimately, tick population size (Knülle and Rudolph 1982, Needham and Teel 1991, Perret et al. 2000, Maldonado-Ruiz et al. 2020). Ticks will begin to desiccate when they are below their critical equilibrium humidity (CEH), which is 80-82% relative humidity for adult *A. americanum* (Hair et al. 1975, Knülle and Rudolph 1982). While desiccation is primarily driven by low relative humidity (at or below CEH), higher temperatures lead to more water molecules escaping the cuticle due to kinetic energy (Sauer and Hair 1986). For *A. americanum,* temperatures in excess of 35 °C can contribute to desiccation and mortality (Sauer and Hair 1971), while all terrestrial arthropods, including ticks, lose their capacity to retain moisture at temperatures in excess of 40 °C (Londt and Whitehead 1972). Relative humidity and temperature can be converted to vapor pressure deficit (VPD) using the formula from Murray (1967). The daily hazard of tick mortality has been shown to increase drastically with increasing levels of daily maximum VPD (e.g., > 2 kPa) of tick species such as *Ixodes scapularis* and *Ixodes ricinus* (Herrmann and Gern 2010, Brunner et al. 2023). Increases in daily maximum VPD has also been associated with a reduction in tick counts and limiting questing activity for *Ixodes* spp. (Sambado et al. 2024) and has shown to be an important predictor of unsuitable habitat for *A. americanum* (Rochlin et al. 2023). Tick abundance, including local abundance of *A. americanum,* can also be regulated by host activity (Paddock and Yabsley 2007, Rochlin et al. 2022). White-tailed deer (*Odocoileus virginianus*) serve as key hosts for all three life stages of *A. americanum* (Bloemer et al. 1986), and are important reservoir hosts for multiple pathogens transmitted by *A. americanum* (Allan et al. 2010b).

### Field experiment: Tick survival and microclimate conditions

To test if ticks survived longer in *Imperata-*invaded areas, we established an experiment in a planted longleaf pine (*Pinus palustris*) stand with fine sandy soils near La Crosse, FL (29.897° N, -82.420° W) that had distinct *Imperata*-invaded and native plant dominated areas (Appendix S1: Figure S2, S3). The invaded area was highly invaded by *Imperata* and spanned approximately 6300 m^2^, while the surrounding area was not invaded and dominated by native plant species. The native plant community was typical of a longleaf pine stand, with *Aristida stricta* (wiregrass), *Rubus flagellaris* (common dewberry), *Carya sp.* (hickory), and *Andropogon* spp. There was no recent chemical or mechanical management in either the invaded or native areas, and the native plant community included no other non-native plant species. Because we conducted our experiment within a forest stand with distinct invaded (*Imperata*) and native areas in close proximity that shared the same overstory structure, we expected any differences in microclimate conditions to be due to invasion, not other underlying factors (see Results).

We established 24 plots (1m x 1m), 12 in invaded habitat and 12 in native plant dominated habitat. Plots were randomly established along two transects, one in each plant community type, with at least 10m spacing. At the center of each plot, a 300-micron 10 cm x 20 cm nylon mesh bag (Skimz, Singapore) containing 10 nymph and 10 adult (5 female and 5 male) *A. americanum* obtained from the National Tick Research and Educational Resource (Department of Entomology and Plant Pathology, Oklahoma State University, Stillwater, OK) was placed in a 50 cm tall cylindrical wire mesh exclosure to deter wildlife interference (Appendix S1: Figure S2). We used lab-reared instead of wild-caught ticks to control for tick age (within life stage) and to standardize the range of tick exposure to environmental conditions prior to the experiment. Each mesh bag containing ticks was oriented vertically to facilitate natural diurnal tick questing (i.e., host seeking behavior) and, although there was no substrate within the bags, the bags remained in contact with the soil surface to allow for ticks to seek refuge below the leaf litter to avoid desiccation stress (Schulze and Jordan 2003, Ostfeld and Brunner 2015) throughout the experiment (Appendix S1: Figure S2).

The start date of the survival experiment (June 21, 2018) coincided with typical off host periods (between blood meals) for *A. americanum* in the region (Davidson et al. 1994, Gleim et al. 2014). During these long non-parasitic phases, ticks seek refuge in areas with microclimates that promote water balance (Knülle and Rudolph 1982, Davidson et al. 1994). Mesh bags were examined every 7-10 days for the first month and then every 10-14 days afterward to determine days survived for both nymph and adult ticks. During each visit, bags were removed from the cages but not opened, and survival status of each individual tick was determined by assessing the response of the tick to CO_2_ stimulation from human breath (Steullet and Guerin 1992). Number of days survived for individual ticks was determined by the time between the start of the survival experiment and the date when the tick was observed not responding to CO_2_ stimulation. Visits continued to the site until May 23, 2019, when all ticks had died.

To assess plant community structure and abiotic conditions in invaded and native plant dominated areas, at the start of the survival experiment we collected understory herbaceous plant height and standing biomass, light availability at the soil surface, and overstory canopy cover at each plot. Plant height was measured as the tallest point of living leaf tissue from soil surface. We collected aboveground biomass by clipping at the soil surface in 25 cm x 25 cm quadrats and dried at 60 °C to a constant mass and weighed (± 0.01g). The 25 cm quadrats were placed adjacent to each 1 m experimental plot, in an area representative of the plant communities in the experimental plots and far enough away (∼1-2m) to not alter microclimate conditions in the experimental plot. Photosynthetically active radiation (PAR, i.e., light) was measured at ground level in each plot using an ACCUPAR LP-80 PAR/LAI ceptometer (Decagon Device Inc., Pullman, WA, USA). To determine if the pine tree overstory was similar in native and invaded areas and was not a confounding factor with invasion, overstory canopy cover was estimated using a concave spherical densiometer (Forestry Suppliers Inc., Jackson, MS, USA) by taking the average of four readings facing each cardinal direction above each plot.

Throughout the experiment we recorded ground-level microclimate conditions (temperature and relative humidity) at five-minute intervals immediately adjacent to each enclosure containing the mesh bags using HOBO U23 Pro v2 loggers (Onset Computer Corporation, Bourne, MA). Each logger was housed in a capped 18-inch length of 3.8 cm diameter PVC pipe to protect it from rainfall and direct sunlight. Holes were drilled on four sides six inches apart along the length of the pipe to allow for airflow and better reflect ambient temperature and relative humidity (see Appendix S1: Figure S2a-b). We used the temperature and relative humidity data at each five-minute interval to calculate VPD (Murray 1967).

### Statistical analysis: Site conditions

We conducted our statistical analysis of the tick survival field experiment in several phases. First, we conducted analyses to determine a) if *Imperata* invaded plant communities had different characteristics that might influence microclimate conditions and b) if *Imperata* invasion altered tick survival over time. We used multivariate analysis of variance (MANOVA) to test the hypothesis that *Imperata* was associated with different plant community characteristics (i.e., vegetation height, standing biomass, light availability at soil surface, and overstory canopy cover). We also plotted microclimate conditions (i.e., lower average daily maximum VPD, higher average daily minimum relative humidity, and lower average daily maximum temperature) over time to test the hypothesis that microclimates differed between native plant dominated and invaded plots. These analyses were done using the “car” package (Fox and Weisberg 2018, R Core Team 2020) in R version 3.6.1.

### Statistical analysis: Tick survival and microclimate conditions

We used linear models using general least squares regression to assess how the number of days survived for individual *A. americanum* of each life stage was related to the plant community and VPD (daily maximum VPD). The general linear mixed effects model on number of days survived for adult or nymph life stage ticks included the fixed effect terms of treatment (native), maximum daily VPD experienced prior to mortality, and their interaction. Maximum daily VPD prior to mortality was recorded as the maximum daily VPD measured at each plot between survival checks (i.e., maximum VPD between the time when the tick was last observed alive and when it was observed as dead). If data loggers failed or malfunctioned, then data from that time interval was omitted from analyses. We included plot as a random effect as well as a compound symmetry covariance structure based on BIC to account for repeated measures over time. Since nymphs are generally more susceptible to desiccation due to their higher surface:volume ratio and lower biomass energy reserves compared to adult ixodid ticks (Knülle and Rudolph 1982, Needham and Teel 1991), we fit models for nymph and adult life stage ticks separately using the “nlme” package (Pinheiro et al. 2025). Degrees of freedom were calculated using “lmertest” (Kuznetsova et al. 2017) in R 3.6.1 (R Core Team 2019).

### Field survey: White-tailed deer activity, tick abundance, and microclimate

To examine the relationship between *Imperata* invasions and white-tailed deer activity and tick abundance, we conducted field surveys in invaded and native dominated plant communities across eight sites in north central Florida (Appendix S1: Figure S3). Invaded plots were selected if the *Imperata* invasion covered a minimum of 500 m^2^, and native plant dominated plots were selected if they had visually similar overstory tree canopy structure to that of the invaded plots. Since we encountered many large invasions with few comparable native plant communities, at some sites we sampled multiple invaded plots per site. We sampled a total of 34 plots, 22 plots in *Imperata* invaded areas and 12 in native plant dominated areas. Due to limited supply of camera traps, a sub-set of the plots were sampled for white-tailed deer activity (28 plots total, 17 in *Imperata* invaded areas and 11 in native plant dominated areas, Appendix S1: Table S1). Plots were also sampled for tick abundance and some plots were sampled multiple times, yielding 51 tick sampling events, 32 in *Imperata* invaded areas and 19 in native plant communities (Appendix S1: Table S2). Corresponding pairs of native and invaded plots that were resampled were chosen haphazardly and revisited after a minimum of three weeks from the previous sampling event (Appendix S1: Table S2). Within a site, invaded and native plant dominated plots were located at least 20 m away from each other. *Imperata* invasions varied in size across sites but covered a minimum of 500 m^2^ to more than 1 ha. We established plots (10 m radius) under similar overstory tree species and canopy cover between invaded and native plots, so the primary difference in habitats was presence or absence of the invader.

We placed two motion-activated cameras (Primos 12MP Proof Cam 02 HD, Primos Hunting, Flora, MS) in each plot to monitor wildlife activity. We focused our analysis on white-tailed deer activity only due to the tall vegetation in invaded areas limiting detection of smaller wildlife hosts with camera traps (Appendix S1: Figure S1). All cameras were set to take a burst of three photos on a 30 second delay, meaning that after the cameras were motion activated there would be a 30 second delay before the cameras were able to be activated again. From the burst of three photos, we analyzed only one photo to count occurrences. In the rare event that an individual or multiple deer were activating the camera by staying in frame for longer than 30 seconds, the deer in these photos were recorded as one occurrence. We made no effort to correct for particular individuals as we were interested in measuring white-tailed deer activity per day rather than number of unique hosts. Cameras were placed simultaneously at native and invaded plots so that deer activity could be calculated across the same time frame. At the end of the experiment photos were examined by three individuals for presence of white-tailed deer (Appendix S1: Table S1). Cameras were positioned on opposite sides of the plot at a distance between 10 m and 15 m from the center of the plot and approximately 1.5 m aboveground (Appendix S1: Figure S4). Cameras were positioned so that their fields of view did not overlap, maximizing the total area of the plot being sampled. Cameras were active from 11 April to 1 June 2019, which coincided with questing activity for nymph and adult *A. americanum* in the region (Davidson et al. 1994, Gleim et al. 2014). In each field site we also recorded ground-level microclimate conditions (temperature and relative humidity) from 13 June to 1 August 2019 at five-minute intervals using HOBO U23 Pro v2 logger and PVC housing as described in the field experiment above.

While the cameras were deployed, we concurrently estimated tick abundance for each plot using carbon dioxide-baited traps (hereafter CO_2_ trap). The CO_2_ trap is one of the most effective means of sampling *A. americanum* nymph and adult life stage ticks (Schulze et al. 1997) and is less biased by different vegetation structures (e.g., a plant invasion) than drag sampling (Kensinger and Allan 2011). The CO_2_ traps consisted of a mini cooler mounted on a sheet of plywood (45 cm x 60 cm) with vent holes drilled along the sides of the cooler near the base to disperse gaseous CO_2_ from sublimating dry ice (Appendix S1: Figure S5). For each sampling event, double-sided carpet tape was placed around the perimeter of the plywood to capture ticks attracted by the CO_2_. Each trap was filled with approximately 2 kg of dry ice and left in the field for 24 hours before ticks were collected. We placed four CO_2_ traps per plot, and traps were placed at least 10m apart from each other within a plot. CO_2_ traps can attract ticks that are within a 5m radius of the trap (Kensinger and Allan 2011). When plots were resampled, traps were placed in new locations that were greater than 5m from the previous sampling location. This sampling approach maximized the area of the 10m radius plot that could be sampled with CO_2_ traps (Appendix S1: Figure S4). In total there were 51 tick sampling events, and with four CO_2_ traps per sample, there were 204 trap nights in the study. Traps were purposefully not placed in plot centers to ensure that new areas of the plot were examined during each sampling event. Ticks ensnared on the tape were counted and preserved in 70% ethanol for identification and to determine life stage.

### Statistical analysis: Tick abundance and white-tailed deer activity

Relative white-tailed deer activity was estimated and standardized by dividing the number of camera trap photos containing deer by the number of days (total hours / 24) the camera was deployed and operating. This approach provided data in units of deer photos/day and allowed for comparison of data from cameras with different lengths of operable deployment. Our approach was intended to detect relative differences in activity by white-tailed deer in invaded and native areas, not to estimate white-tailed deer population sizes.

We fit a negative binomial general linear mixed model to examine the relationship between white-tailed deer activity and concurrent estimates of tick abundance from 11 April to 1 June 2019 in native and invaded plant communities. We fit a model of tick abundance (nymph + adult ticks) using plant invasion status (native vs. invaded), white-tailed deer activity, and their interaction, as fixed-effect terms and site as a random effect term to account for repeated measures. Tick abundance includes both nymph and adult life stage ticks, because white-tailed deer serve as an important host for both life stages of *A. americanum* (Bloemer et al. 1986). Models were fit using the “lme4” package in R 3.6.1 (Bates et al. 2014, Team 2019). Two sided t-test was also used to test for differences in relative host activity and in tick abundance between native and invaded plant communities.

## Results

### Field experiment: Site conditions

There was a significant difference between the native and invaded plant community types on the combined dependent variables average plant height, plant biomass, light availability at the soil surface, and overstory canopy cover (MANOVA; Pillai’s Trace _1,22_ = 0.922; *p* < 0.0001). In separate one-way ANOVAs, average plant height was 268% higher (*F*_1,22_ = 179.83; *p* <0.001) and plant biomass was 408% greater in invaded than native areas (*F*_1,22_ = 29.29; *p* <0.001) (Appendix S1: Figure S6a, b), which corresponded to 91% less light availability at the soil surface in invaded areas (*F*_1,22_ = 21.22; *p* = <0.001) (Appendix S1: Figure S6c). However, there was no difference in overstory canopy cover in invaded and native areas (*F*_1,22_ = 0.15; *p* = 0.706), indicating differences in plant communities and understory light availability were likely driven by presence of *Imperata* and not differences in overstory canopy cover (Appendix S1: Figure S6d).

### Field experiment: Microclimate conditions and tick survival

Throughout the survival experiment, average daily maximum VPD was approximately 70% lower in invaded (mean ± SE: 0.99 ± 0.04 kPa) compared to native (1.7 ± 0.07, Figure 2a) areas. Average daily minimum relative humidity was approximately 16% greater in invaded (mean ± SE: 79.21% ± 0.79) compared to native (67.98% ± 0.87) (Appendix S1: Figure S7a). Average daily maximum temperature was approximately 13% lower in invaded (mean ± SE: 27.20°C ± 0.34) compared to native (31.29 °C ± 0.40) (Appendix S1: Figure S7b).

**Figure 2.**
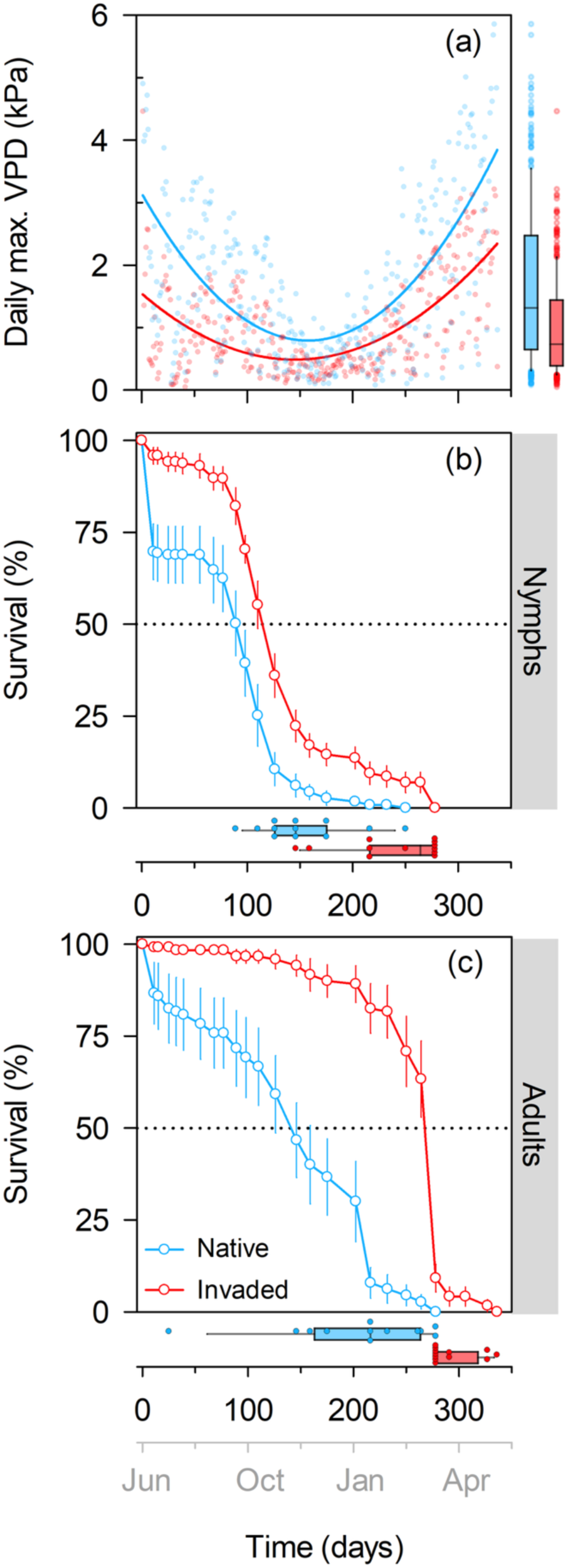
Microclimate conditions and tick survival per plot in native and invaded areas. Average daily maximum vapor pressure deficit (VPD, kPa) (a), points are the average of maximum daily VPD across all plots in each plant community type (n = 12). Solid lines in panel (a) indicate best fit quadratic polynomials for native (blue) and invaded (red) areas. Box plots to the right of panel (a) indicate the median and 25^th^ and 75^th^ percentile, with 10^th^ and 90^th^ percentile whiskers for daily maximum VPD (kPa). Percent survival (mean ± SE) of (b) nymph and (c) adult *Amblyomma americanum* in native (blue) and invaded (red) plots over time; the second x-axis shows the month of the year. Box plots below panels (b) and (c) are distributions of time to 100% mortality across the 12 plots showing the median and 25^th^ and 75^th^ percentile, with 10^th^ and 90^th^ percentile whiskers.

Regardless of sex or life stage, ticks survived longer in invaded compared to native plant communities (Figure 2 c, d). Time to 100% mortality per plot was, on average, 57% longer for nymphs and 45% longer for adults in the invaded compared to native dominated plots (Figure 2 c, d). Mean ± SE nymph survival was 130 ± 5.4 days in invaded areas versus 80.5 ± 5.0 days in native vegetation, while mean adult survival was 256 ± 4.8 and 140 ± 7.3 days in invaded and native areas, respectively. For nymph life stages, daily maximum VPD (kPa) was significantly related to fewer days survived in both native and invaded plant communities (Figure 3a) (Appendix S1: Table S3). Daily maximum VPD (kPa) was significantly related to fewer days survived for adult life stage ticks in the native plant dominated community (Figure 3b) (Appendix S1: Table S3). However, in invaded areas there was a significant, positive relationship between daily maximum VPD and days survived (Figure 3b) (Appendix S1: Table S3).

**Figure 3.**
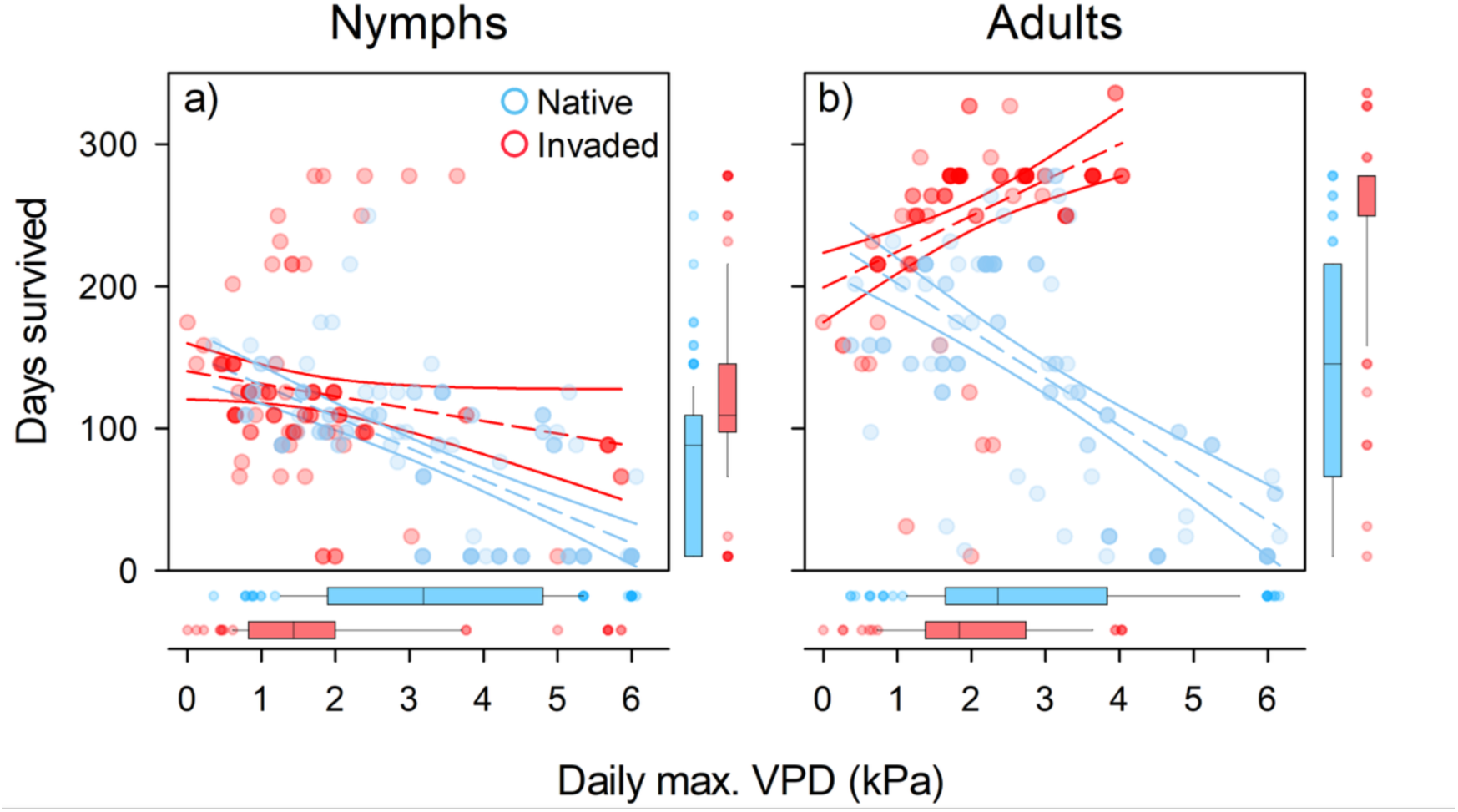
Number of days survived for individual nymph (a) and adult (b) life stage ticks and the daily maximum VPD experienced in native (blue) and invaded (red) plots between survival checks (the time interval when the tick was last seen alive and when it was recorded dead). Best fit lines (dashed) with 95% confidence interval (solid) are from linear mixed effects models. Box plots indicate median and 25th and 75th percentile, with 10th and 90th percentile whiskers for days survived (y-axes) and daily maximum VPD experienced by individual ticks prior to mortality in each treatment (x-axes).

### Field survey: Microclimate, white-tailed deer activity, and tick abundance

Across the eight field sites, microclimate conditions (vapor pressure deficit, relative humidity and maximum temperature) in invaded and native plant communities were similar to what was observed at the field experiment site, though over a much shorter time period (Appendix S1: Figure S8). Mean ± SE of daily maximum VPD (kPa) was 0.4 ± 0.03 in invaded areas and 1.8 ± 0.07 in native areas, minimum relative humidity (%) was 91.4 ± 0.7 in invaded areas and 67.5 ± 1.0 in native areas, while average daily maximum temperature (°C) was 30.0 ± 0.1 and 34.00 ± 0.2 in invaded and native areas, respectfully (Appendix S1: Figure S8).

There were no differences in relative white-tailed deer activity between native and invaded plots (*t_44_* = 1.01, *p* = 0.318) (Appendix S1: Figure S9). There were also no differences in average tick abundance (nymph [*t_43_* = -0.713, *p* = 0.480]; adult [*t_43_* = -1.031, *p* = 0.308] life stage ticks) between native and invaded areas (Appendix S1: Figure S10). In native communities, tick abundance increased with white-tailed deer activity (deer photos/day) (estimate = 0.6 ticks per deer photo/day; 95% CI = 0.1 – 1.1) (Figure 4, Appendix S1: Table S4), while in invaded habitats, white-tailed deer activity was not related to tick abundance (estimate = -0.4 ticks per deer photo/day; 95% CI = -0.9 – 0.1). For plots where no white-tailed deer were detected (i.e., intercept term), invaded plant communities had positive tick abundances (estimate = 1.4 ticks; 95% CI = 0.2 – 2.5) (Figure 4, Appendix S1: Table S4). There was one plot with both the highest tick abundance and highest deer activity that could have strongly influenced the results, so we ran the analysis using the same linear mixed effects model but with that data point removed. Omitting that one data point resulted in no association between white-tailed deer activity and tick abundance (estimate = -0.4 ticks per deer photo/day; 95% CI = -0.9 – 0.1) (Appendix S1: Table S5), but there was a significant intercept term (estimate = 1.3 ticks; 95% CI = 0.1 – 2.6) (Appendix S1:Table S5), indicating that tick abundance was greater in invaded plant communities even when no white-tailed deer were detected.

**Figure 4.**
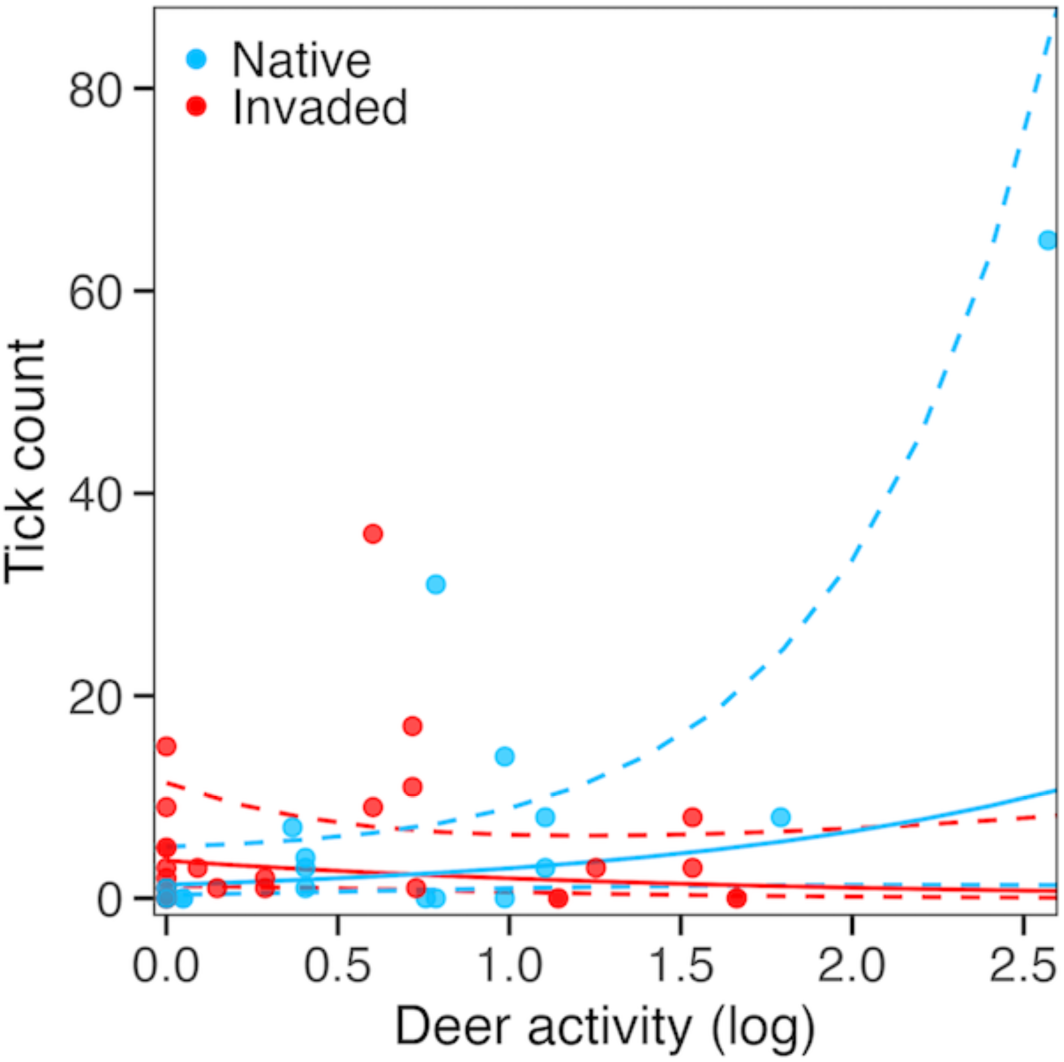
Relationship between deer activity (deer photos/day) and tick abundance (nymph and adult ticks per plot) in native (blue) and invaded (red) plant communities. Best fit lines (solid) from linear mixed effect models with 95% confidence interval (dashed).

## Discussion

These results contribute to a growing understanding that introductions of non-native species may have a myriad of indirect effects on ecological communities (Pejchar and Mooney 2009, Linders et al. 2019). We documented that the invasive plant species cogongrass was associated with greater tick survival time, which could maintain tick abundances even though invaded areas may have less activity by the common host white-tailed deer. Consistent with our hypothesis about differences in microclimate due to invasion, there was greatly prolonged tick survival in invaded areas, which had lower vapor pressure deficit corresponding to both lower temperatures and higher humidity. We had also hypothesized that invaded areas would have greater tick abundance than native-dominated areas because of longer tick survival times and greater white-tailed deer activity. Contrary to this hypothesis, we observed no overall difference in white-tailed deer activity between native and invaded areas and detected no difference in average tick abundance between the two areas during the time period of peak tick abundance (Davidson et al. 1994, Gleim et al. 2014). There was a weak positive relationship between tick abundance and white-tailed deer activity, though only in native-dominated areas, and the pattern was driven by one of the eight sites (Figure 4). Together, these results suggest the invasive plant is influencing multiple, potentially counteracting, mechanistic pathways contributing to tick exposure risk.

Our results show that tick abundances in *Imperata-*invaded plant communities may be maintained even with relatively low white-tailed deer activity due to more favorable microclimate conditions that prolong off-host tick survival. Exploration of the relationship between tick mortality and VPD revealed that survival of nymphal ticks in both vegetation types and adult ticks in native vegetation declined with increasing VPD. Conversely, adult tick survival in invaded habitats was prolonged by increasing VPD. We posit that this counterintuitive result stems from the physiological buffering provided by *Imperata*, where invaded sites rarely exhibited conditions that have been shown to produce rapid mortality for other tick species in laboratory conditions (i.e., VPD > 2 kPa) (Herrmann and Gern 2010). Instead, adult ticks in invaded habitats lived long enough that they eventually died as a result of loss of metabolic reserves (Alasmari and Wall 2021) or other stressors that were not measured. More generally, invasive plant buffering of microclimate conditions that would otherwise be stressful for ticks suggests that invasions may facilitate tick survival under the hotter and drier conditions expected with climate change (McDowell et al. 2022).

Deer are widely considered a key contributing factor to abundances of several North American tick species but the shape of this relationship is often difficult to detect (Kugeler et al. 2016, Hofmeester et al. 2017). Other studies have emphasized the complexity of this relationship, detecting either a weak positive relationship (Ostfeld et al. 2018) or no association between deer abundance and tick abundance (Hofmeester et al. 2017). We essentially found the same pattern: in our analysis of data from all eight field sites, we detected a weak positive relationship between deer and tick abundance in native plant communities but no relationship when an influential site was removed (Figure 4). In contrast to previous studies (Williams et al. 2009, Allan et al. 2010a), we did not find a relationship between tick abundance and deer activity in invaded plant communities; however, both of those studies evaluated non-native invasive shrubs, which may promote deer activity by providing forage or cover. Although other species are known to use *Imperata* invaded areas for refuge, e.g., wild turkey (*Meleagris gallopavo,* Cabrera et al. 2021) and cotton rats (*Sigmodon hispidus*, Borden et al. 2022), we found no difference in abundance of white-tailed deer based on camera trap detections in *Imperata* invasions compared to native plant communities.

Climate change is predicted to facilitate the spread of *Imperata,* expanding its range from southeastern Gulf States to as far north as Virginia along the eastern USA seaboard (Bradley et al. 2010). As *Imperata* spreads northward, it is likely to interact with other tick species, including the blacklegged tick *Ixodes scapularis,* the primary vector for Lyme disease in the United States, which could increase tick-borne disease risk (Steere et al. 2004). The geographic distribution of the black-legged tick is also expanding, with an increase in the number of USA counties reporting its presence from 1996-2005 (Eisen et al. 2016) facilitated by ongoing landscape change (Gardner et al. 2020). The blacklegged tick has similar physiological thresholds to *A. americanum* (Berger et al. 2014), suggesting it too may benefit from the low vapor pressure deficit generated by *Imperata* invasions.

This study uniquely demonstrated how differences in abiotic conditions in invaded areas can be associated with differences in tick abundance and thus disease exposure risk. Despite the duration and precision of the survival assay and the scale of the field study, there were limitations. For example, although we documented much longer tick survival in invaded areas, and evidence for microclimate conditions driving that pattern, additional studies replicated across a larger environmental gradient of invasion would be useful for validating this result. In addition, including sites that vary in invader density or biomass could inform management of highly invaded areas to reduce tick-borne exposure risk. While we found evidence for microclimate driven survival, we found less support for increases in host habitat use within invaded plant communities. We focused on white-tailed deer activity, in part because of invader height and difficulty camera trapping rodents or other hosts with our methods; however, quantifying other hosts (small- and medium-sized mammals, birds, etc.) in invaded areas that are known to influence tick-borne disease risk could resolve the relative influence of host abundance and microclimate-driven survival on tick abundance (Greiner et al. 1984, Clark 2004, Ostfeld et al. 2018). Additionally, since CO_2_ traps only capture host-seeking ticks, actual tick abundance could be greater in invaded areas if *Imperata* improves microclimate to the point where ticks might be more selective about when they expend energy to pursue a host.

The dense herbaceous structure of *Imperata* invasions differ drastically from that of the woody and grassy invaders that have been studied previously, making comparisons across species and systems difficult. For example, studies evaluating the effects of woody invasive plants on tick abundance (*Lonicera maackii*, *Berberis thunbergia, Rosa multiflora*) in the Midwest or Northeast US (Williams et al. 2009, Allan et al. 2010a, Adalsteinsson et al. 2016) have found that woody invasive plants alter tick abundance predominately by providing host habitat rather than creating more favorable microclimates for tick survival (but see Williams and Ward 2010). By contrast, tick survival assays conducted using the invasive grass *Microstegium vimineum* found that the invader created less favorable microclimate conditions and reduced survival time for off-host ticks (Civitello et al. 2008). Impacts of plant invasions on tick abundance will likely differ by location and species studied, thus there is a need to address how invasive plants impact tick abundance through changes to host activity as well as how tick survival is altered by invader-driven changes in microclimate.

Consistent with Wei et al. (2020), our findings suggest that invaded plant communities play an important, but relatively underappreciated, indirect role in altering human vector-borne disease risk. Specifically, we provide evidence for how tick exposure risk may be driven by different mechanistic pathways in invaded and native plant communities. Invasive plant species have a wide variety of traits that could differentially affect off-host tick survival and host activity. While measurement of microclimate factors and tick survival are straightforward for field ecology research studies, measurement of activity of large-bodied mammals with large home range sizes, such as white-tailed deer, is notoriously difficult. Thus, we encourage future work to explore these relationships among invasive plants, environmental conditions, ticks, and their hosts more generally across other invasive plant and tick species, and under different climates, landscape conditions, and disturbance regimes. Understanding the relative contribution of host use and microclimate across habitats invaded by different species will aid in our ability to determine current, and forecast future, impacts of plant invasions on tick-borne disease risk. More broadly, these findings highlight the complex and indirect mechanisms by which invasive species may impact species interactions.

## Acknowledgements

We thank Taylor Clark for assistance with data collection, Tempest McCabe for feedback and assistance with statistical analysis, and members of the Flory Lab at the University of Florida for comments and feedback on figures and text.

## Funding

This research was funded by the U.S. Strategic Environmental Research and Development Program (RC-2636).

The authors declare no conflict of interest

## Notes

### Competing Interest Statement

The authors have declared no competing interest.

### Summary of Updates

Updated analyses to include VPD instead of RH and temperature. Significant updates to the introduction and discussion.

