## Supplemental materials for "Mechanistic pathways of tick exposure risk in native and invaded plant communities"

Journal: Ecology

**Supplemental Materials**

**Appendix S1**

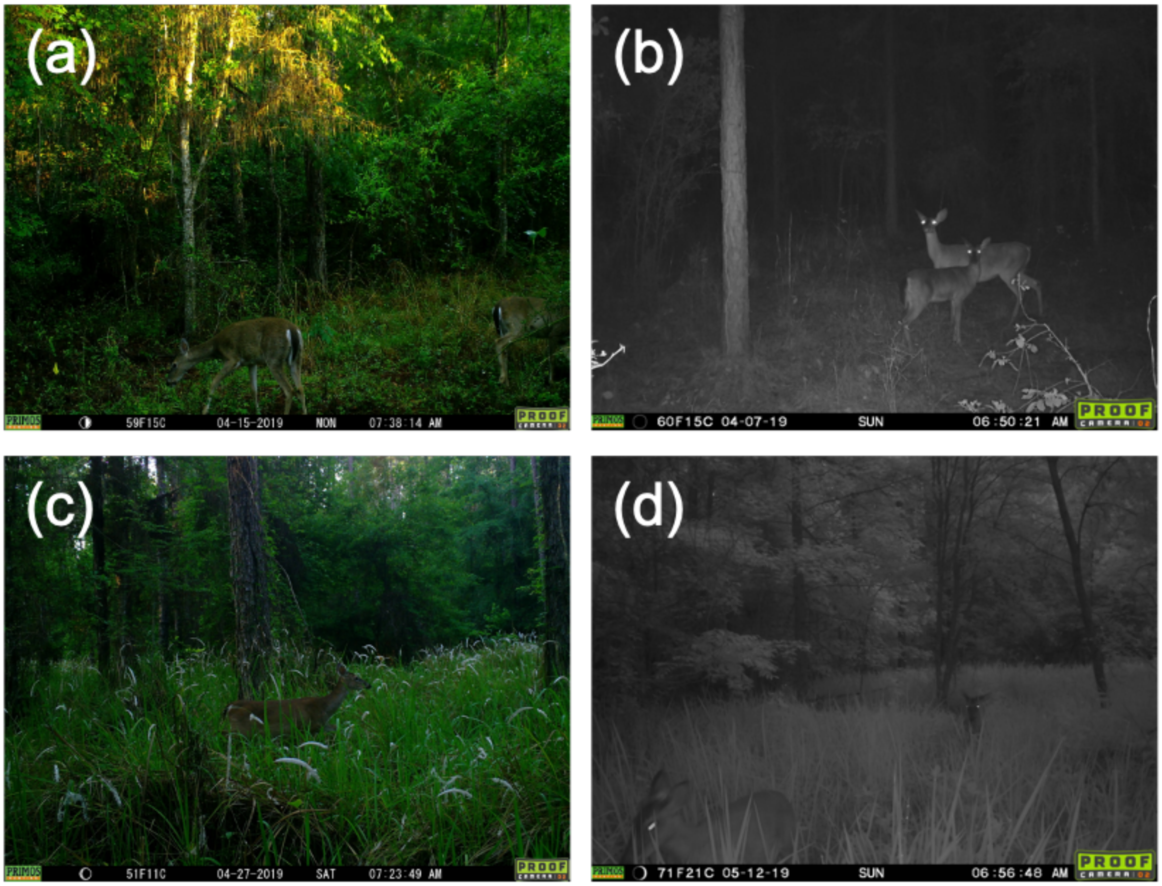

**Figure S1.** Example of White-tailed deer in native (a, b) and *Imperata* invaded plant communities (c, d). Photos taken by Primos 12MP Proof Cam 02 HD, (Primos Hunting, Flora, MS), photograph owner Drew Hiatt.

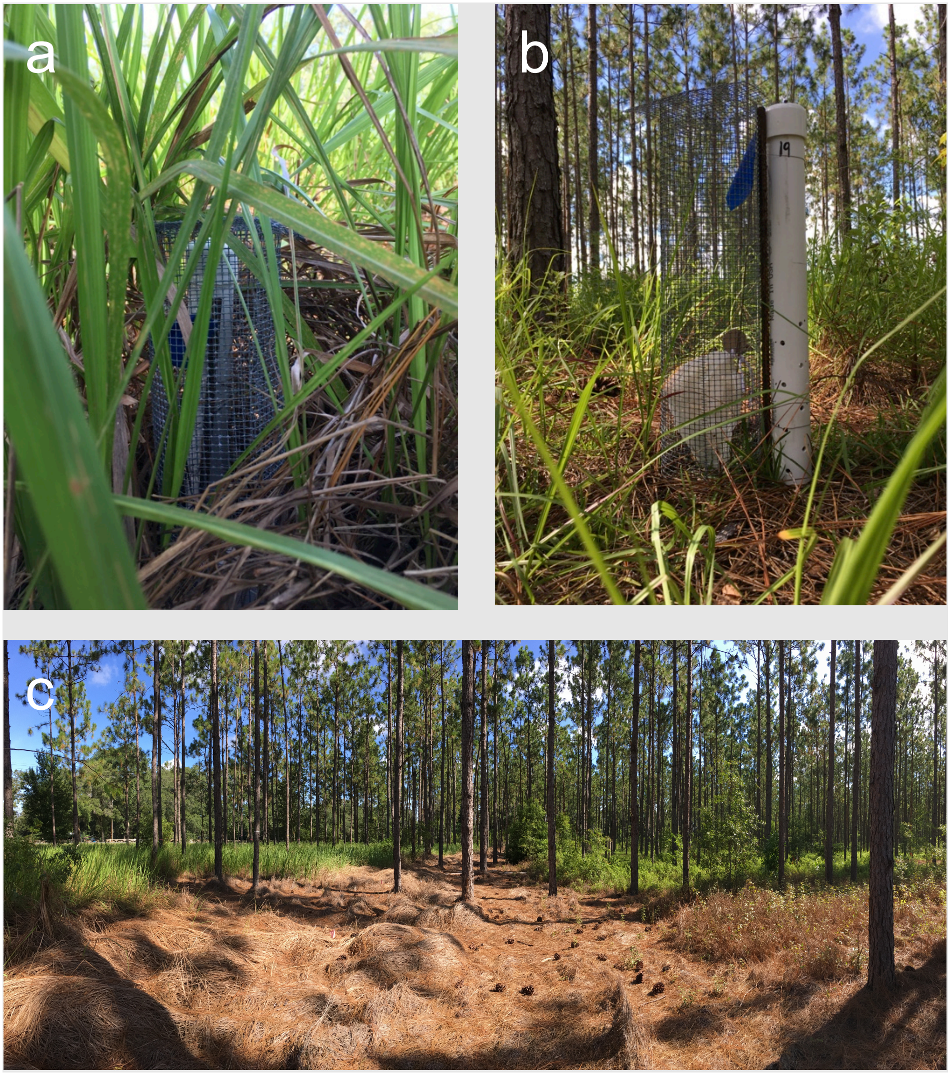

**Figure S2.** Ticks in mesh bags inside wire mesh exclosures along with PVC housing for temperature and relative humidity loggers in (a) invaded and (b) native plant communities. Representation of (c) the field study site showing invaded to the left and native plant communities to the right, separated by managed boundary area to maintain distinct plant communities. Photos taken by Drew Hiatt.

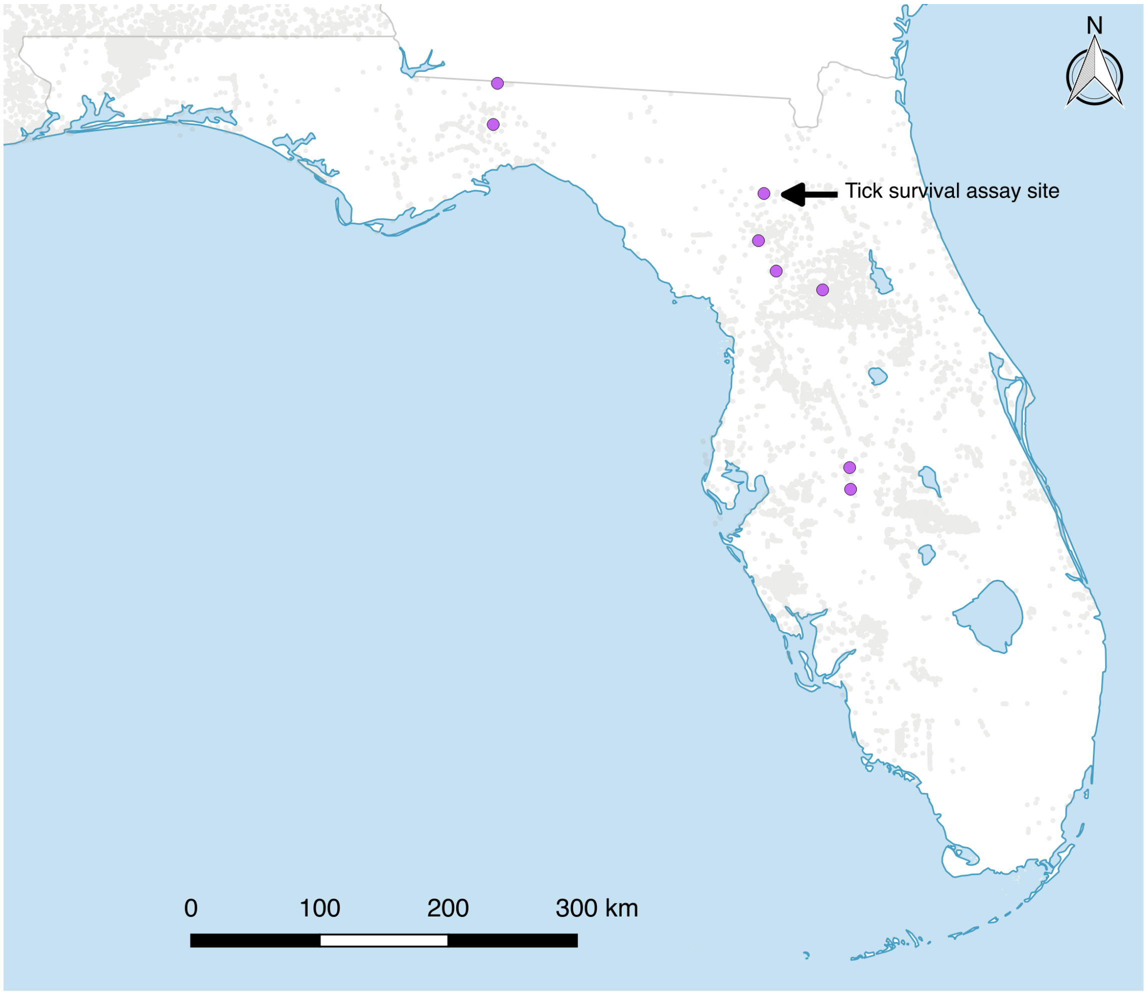

**Figure S3.** Site locations (purple circles) where tick trapping and camera trapping were conducted, site where the tick survival assay is also depicted, background shading indicates documented invasions of cogongrass (n = 74,785) from 1990-2020 (eddmaps.org).

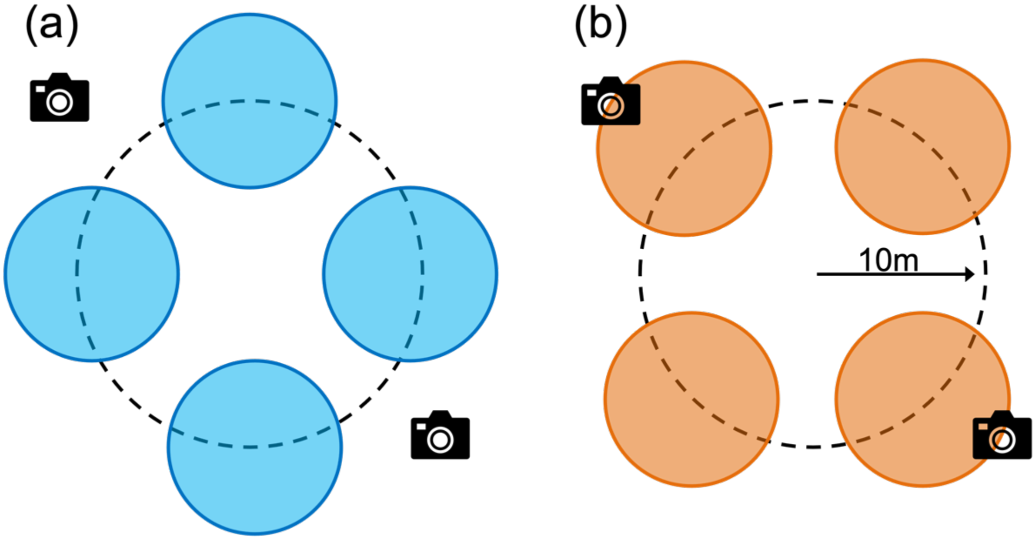

**Figure S4.** An example of one 10m radius plot initially sampled (a) and resampled (b). Filled circles show the sphere of influence for $CO_2$ tick traps (5m radius) for the initial sample (blue) and moved to new locations for resample (orange). Camera icons indicate approximate location of motion-activated cameras. Vector graphics are copyright-free images obtained from stock.adobe.com via the University of Florida's enterprise license

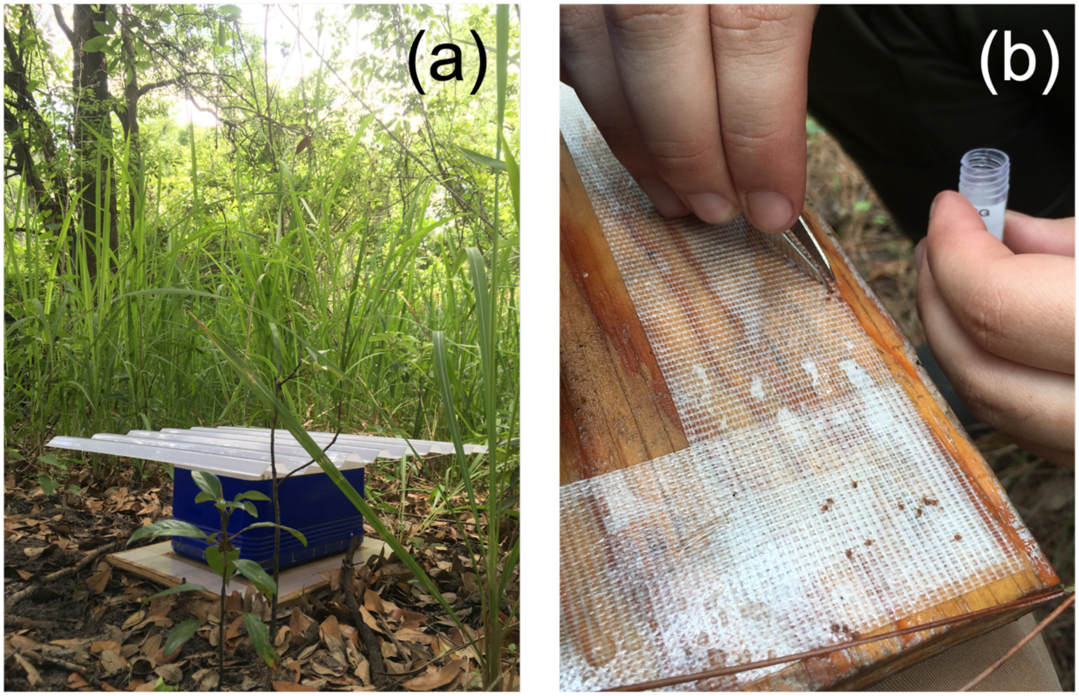

**Figure S5.** $CO_2$ tick trap within an invaded area, equipped with rain shield (a) and ticks being collected from double sided tape on the $CO_2$ trap after 24hrs. Photos taken by Drew Hiatt.

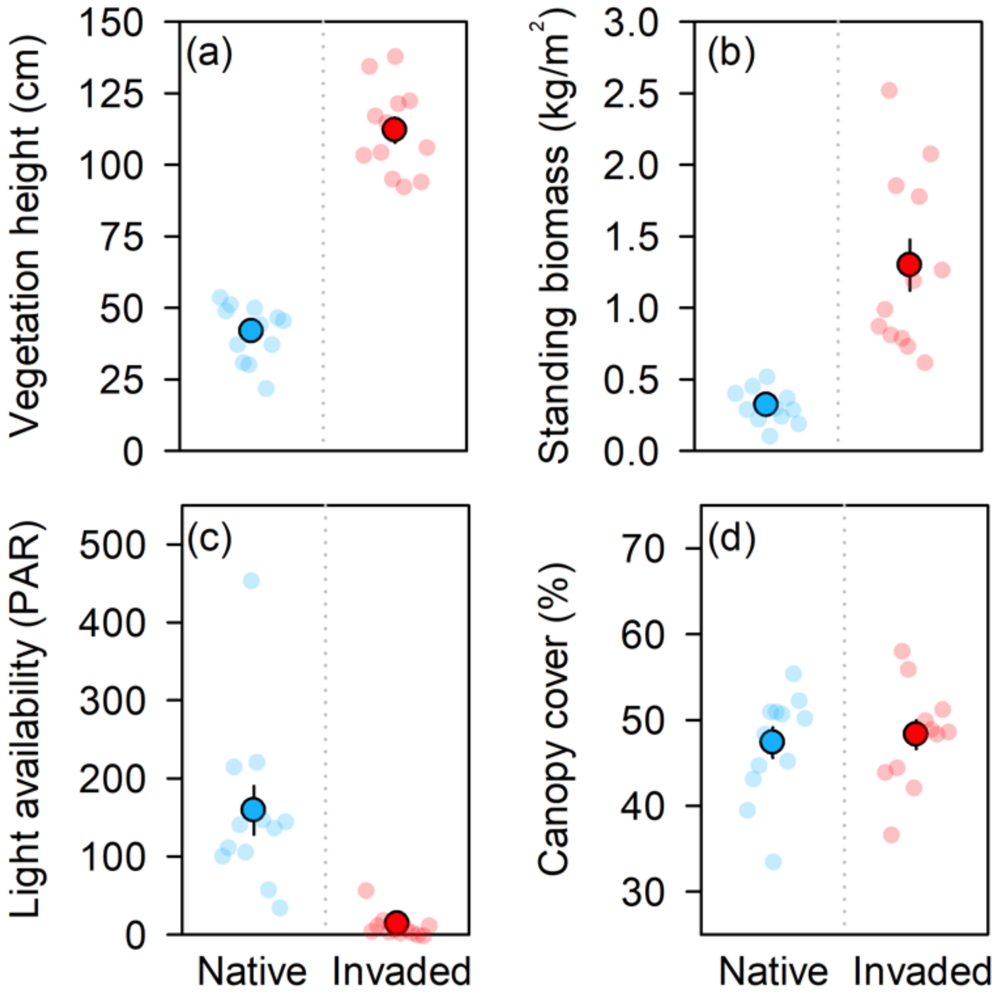

**Figure S6.** Differences (mean ± SE) between native and *Imperata*-invaded plant communities. Plant community structure. (a) vegetation height, (b) standing biomass (kg $m^{-2}$) (c) light availability at the soil surface (μ-moles $m^{-2}$ $sec^{-1}$), and (d) overstory canopy cover. Data for invaded and native plots of the tick survival experiment.

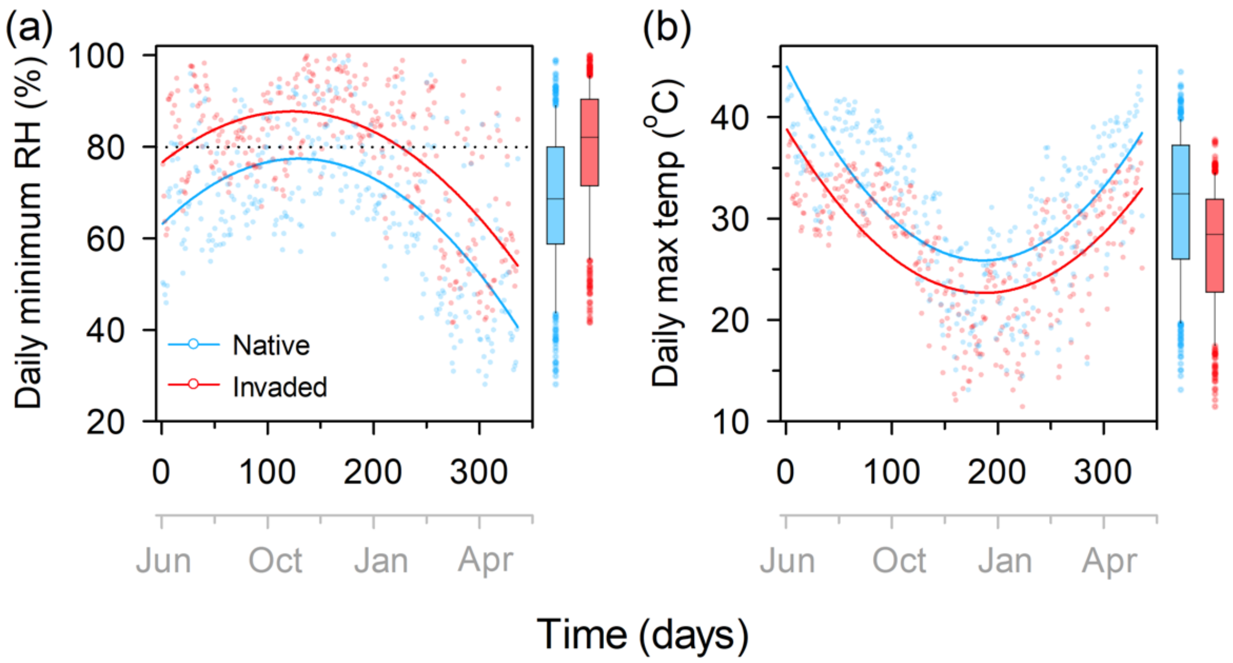

**Figure S7.** Microclimate conditions per plot in native (blue) and invaded (red) areas. Average daily minimum relative humidity (%) (a) and average daily maximum temperature (ºC) (b). Points in (a) and (b) are the daily average across all plots (n = 12) in each treatment. Curved lines indicate are best fit quadratic polynomials for native and invaded areas. Box plots to the right of each panel indicate the median and $25^{th}$ and $75^{th}$ percentile, with $10^{th}$ and $90^{th}$ percentile whiskers for daily minimum RH (%) and daily max temperature (ºC).

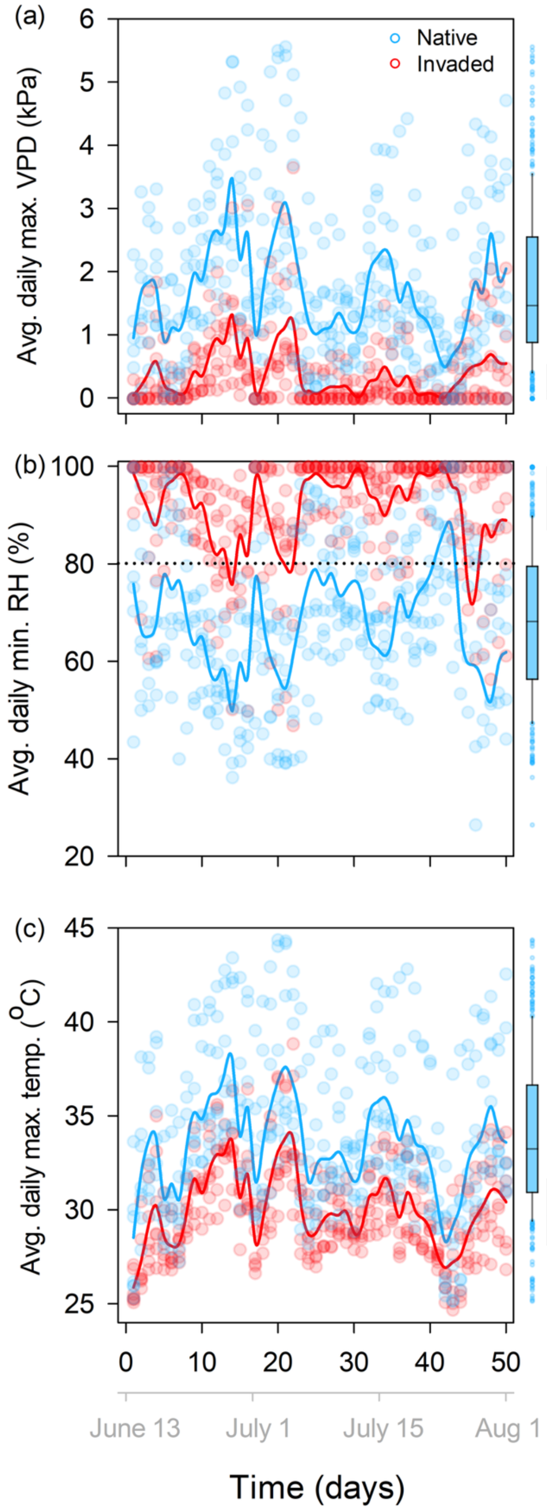

**Figure S8.** Microclimate conditions across regional field sites from (June 13 - August 1) for (a) average daily maximum VPD (kPa), (b) average daily minimum relative humidity

(RH) (%), dotted line indicates critical equilibrium humidity (80% RH) for *Amblyomma americanum* (Hair et al. 1975), and (c) average daily maximum temperature (ºC). Each point represents a data logger (n = 7) in each plant community, solid lines are daily averages across all data loggers in each plant community type for native (blue) and invaded (red) areas. Box plots to the right of each panel indicate the median and 25th and 75th percentile, with 10th and 90th percentile whiskers for each microclimate variable.

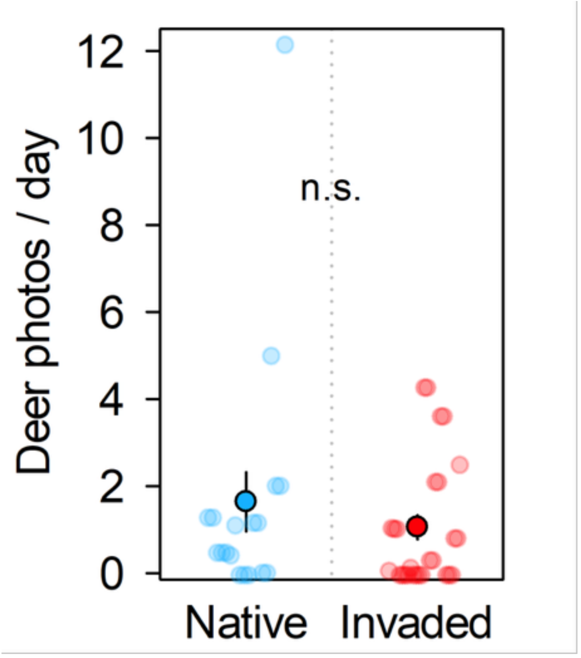

**Figure S9.** Mean ± SE deer activity (deer photos/ day) per plot for field surveys in native and invaded areas.

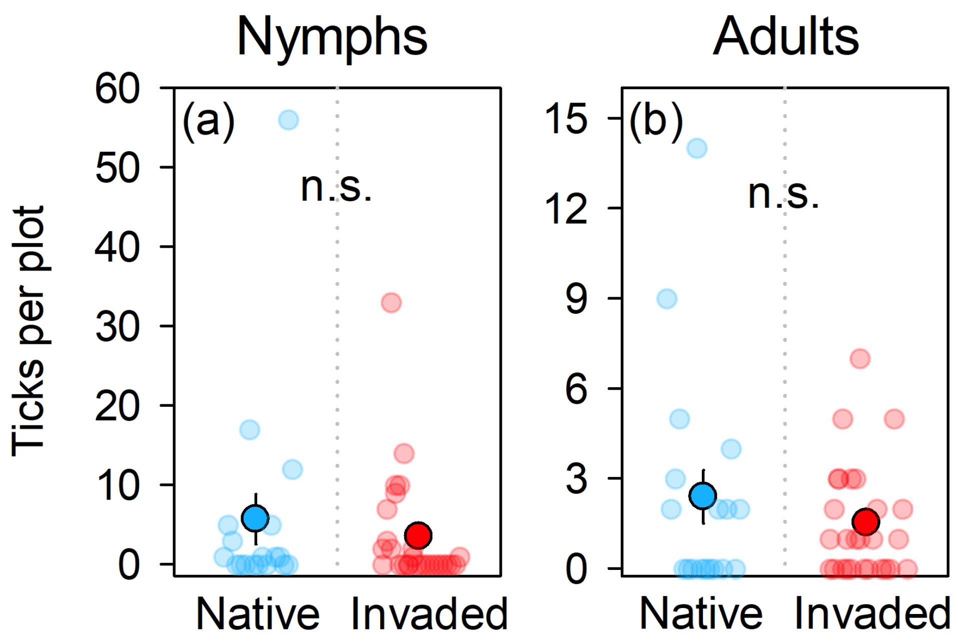

**Figure S10.** Mean ± SE nymph (a) and adult (b) tick abundance per plot for field surveys in native and invaded areas.

**Table S1.** Camera trap data, deer photos, days motion cameras were active, and deer photos per day across plots in native and invaded plots

| Site | Plant comm. | Plot | Deer photos | Days active | Deer photos per day |
|---|---|---|---|---|---|
| *Brown* | Native | n-1 | 290 | 24 | 12.08 |
| | | n-2 | 105 | 52 | 2.02 |
| | | n-3 | 40 | 8 | 5 |
| | Invaded | i-1 | 60 | 24 | 2.5 |
| | | i-2 | 43 | 52 | 0.83 |
| | | i-3 | 0 | 8 | 0 |
| *Hancock* | Native | n-1 | 1 | 22 | 0.05 |
| | | n-2 | NA | NA | NA |
| | Invaded | i-1 | 0 | 42 | 0 |
| | | i-2 | 0 | 20 | 0 |
| | | i-3 | NA | NA | NA |
| | | i-4 | NA | NA | NA |
| *Munson* | Native | n-1 | 43 | 36 | 1.19 |
| | Invaded | i-1 | 131 | 36 | 3.64 |
| *Peace River* | Native | n-1 | 0 | 37 | 0 |
| | Invaded | i-1 | 79 | 37 | 2.14 |
| | | i-2 | 18 | 54 | 0.33 |
| | | i-3 | 124 | 29 | 4.26 |
| | | i-4 | NA | NA | NA |
| | | i-5 | NA | NA | NA |
| *Rivers Edge* | Native | n-1 | 0 | 3 | 0 |
| | Invaded | i-1 | 0 | 1 | 0 |
| | | i-2 | 0 | 3 | 0 |
| *Seay* | Native | n-1 | 78 | 69 | 1.13 |
| | | n-2 | 8 | 18 | 0.44 |
| | Invaded | i-1 | 11 | 69 | 0.16 |
| | | i-2 | 0 | 18 | 0 |
| *Silver Springs* | Native | n-1 | 20 | 40 | 0.5 |
| | Invaded | i-1 | 0 | 40 | 0 |
| *Wes* | Native | n-1 | 106 | 63 | 1.68 |
| | Invaded | i-1 | 42 | 40 | 1.05 |
| | | i-2 | 45 | 42 | 1.07 |
| | | i-3 | 6 | 63 | 0.10 |

**Table S2.** Number of adult and nymph ticks trapped as well as camera trap effort across sites and plots within sites.

| Site | Plant comm. | Plot | Date | Adult | Nymphs | Total | Camera trap data |
|---|---|---|---|---|---|---|---|
| *Brown* | Native | n-1 | 4/5 | 9 | 56 | 65 | Yes |
| | | n-2 | 4/26 | 2 | 1 | 3 | Yes |
| | | n-2 | 5/17 | 3 | 5 | 8 | Yes |
| | | n-3 | 5/17 | 5 | 3 | 8 | Yes |
| | Invaded | i-1 | 4/5 | 1 | 2 | 3 | Yes |
| | | i-2 | 4/26 | 3 | 33 | 36 | Yes |
| | | i-2 | 5/17 | 2 | 7 | 9 | Yes |
| | | i-3 | 5/17 | 5 | 10 | 15 | Yes |
| *Hancock* | Native | n-1 | 4/18 | 0 | 0 | 0 | Yes |
| | | n-1 | 5/8 | 0 | 0 | 0 | Yes |
| | | n-2 | 5/8 | 0 | 0 | 0 | No |
| | Invaded | i-1 | 4/18 | 0 | 0 | 0 | Yes |
| | | i-2 | 4/18 | 0 | 0 | 0 | Yes |
| | | i-3 | 5/8 | 0 | 0 | 0 | No |
| | | i-4 | 5/8 | 1 | 0 | 1 | No |
| *Munson* | Native | n-1 | 5/1 | 14 | 17 | 31 | Yes |
| | | n-1 | 6/7 | 0 | 0 | 0 | Yes |
| | Invaded | i-1 | 5/1 | 7 | 1 | 8 | Yes |
| | | i-1 | 6/7 | 3 | 0 | 3 | Yes |
| *Peace River* | Native | n-1 | 4/11 | 0 | 0 | 0 | Yes |
| | Invaded | i-1 | 4/11 | 0 | 0 | 0 | Yes |
| | | i-1 | 4/18 | 0 | 0 | 0 | Yes |
| | | i-2 | 4/11 | 1 | 0 | 1 | Yes |
| | | i-2 | 4/18 | 2 | 0 | 2 | Yes |
| | | i-3 | 4/11 | 0 | 0 | 0 | Yes |
| | | i-3 | 4/18 | 0 | 0 | 0 | Yes |
| | | i-4 | 5/8 | 0 | 0 | 0 | No |
| | | i-5 | 5/8 | 0 | 0 | 0 | No |
| *Rivers Edge* | Native | n-1 | 5/1 | 0 | 1 | 1 | Yes |
| | | n-1 | 6/7 | 0 | 0 | 0 | Yes |
| | Invaded | i-1 | 5/1 | 5 | 0 | 5 | Yes |
| | | i-1 | 6/7 | 0 | 0 | 0 | Yes |
| | | i-2 | 5/1 | 1 | 0 | 1 | Yes |
| | | i-2 | 6/7 | 2 | 0 | 2 | Yes |
| | | i-3 | 5/1 | 0 | 0 | 0 | No |

**Table S2.** Cont.

| Site | Plant comm. | Plot | Date | Adult | Nymphs | Total | Camera trap data |
|---|---|---|---|---|---|---|---|
| *Seay* | Native | n-1 | 4/24 | 0 | 0 | 0 | Yes |
| | | n-2 | 5/15 | 2 | 5 | 7 | Yes |
| | Invaded | i-1 | 4/24 | 0 | 1 | 1 | Yes |
| | | i-2 | 5/15 | 0 | 0 | 0 | Yes |
| *Silver Springs* | Native | n-1 | 4/5 | 4 | 0 | 4 | Yes |
| | | n-1 | 4/26 | 0 | 1 | 1 | Yes |
| | | n-1 | 5/17 | 2 | 1 | 3 | Yes |
| | Invaded | i-1 | 4/5 | 3 | 2 | 5 | Yes |
| | | i-1 | 4/26 | 0 | 3 | 3 | Yes |
| | | i-1 | 5/17 | 0 | 9 | 9 | Yes |
| *Wes* | Native | n-1 | 4/24 | 0 | 0 | 0 | Yes |
| | | n-1 | 5/15 | 2 | 12 | 14 | Yes |
| | Invaded | i-1 | 4/3 | 3 | 14 | 17 | Yes |
| | | i-1 | 5/15 | 1 | 10 | 11 | Yes |
| | | i-2 | 4/24 | 1 | 0 | 1 | Yes |
| | | i-3 | 5/15 | 1 | 2 | 3 | Yes |

**Table S3(new).** Tick survival (days) and daily maximum VPD (Figure 3 a & b)

| | Term | Estimate | 95% CI | t value | Df | p value |
|---|---|---|---|---|---|---|
| **Nymphs** | Intercept | 150.57 | (127.8, 173.4) | 13.02 | 185 | **<0.0001** |
| | Treatment (native) | 17.46 | (-18.2, 53.1) | 0.96 | 22 | 0.335 |
| | Max VPD | -13.09 | (-21.89, -4.3) | -2.94 | 185 | **0.037** |
| | Trt (native)*Max VPD | -13.69 | (-24.8, -2.6) | -2.44 | 185 | **0.016** |
| | Random effect | | | | | |
| | Plot ID: Rho = 0.24 | | | | | |
| | $R^2$: 0.333 | | | | | |
| **Adults** | Intercept | 184.8 | (148.5, 221.2) | 10.02 | 188 | **<0.0001** |
| | Treatment (native) | 24.8 | (-24.9, 74.5) | 0.98 | 20 | 0.326 |
| | Max VPD | 32.22 | (20.5, 43.9) | 5.42 | 188 | **<0.0001** |
| | Trt (native)*Max VPD | -56.98 | (-71.2, -42.8) | -7.92 | 188 | **<0.0001** |
| | Random effect | | | | | |
| | Plot ID: Rho = 0.49 | | | | | |
| | $R^2$: 0.591 | | | | | |

**Table S4.** Negative binomial linear regression models for habitat type, host activity, and their interaction on tick abundance from field observations (Figure 4)

| | **Term** | Estimate | Std. Error | z value | df | p value |
|---|---|---|---|---|---|---|
| **Tick abund.** | Intercept | 1.35 | 0.60 | 2.24 | 45 | **0.025** |
| | Treatment (native) | -0.81 | 0.51 | -1.59 | 45 | 0.111 |
| | Deer activity | -0.42 | 0.25 | -1.67 | 45 | 0.093 |
| | Trt (native)*Deer activity | 0.59 | 0.26 | 2.23 | 45 | **0.026** |

**Table S5.** Negative binomial linear regression models for habitat type, host activity, and their interaction on tick abundance from field observations *with data point omitted.*

| | **Term** | Estimate | Std. Error | z value | df | p value |
|---|---|---|---|---|---|---|
| **Tick abund.** | Intercept | 1.34 | 0.63 | 2.14 | 44 | **0.033** |
| | Treatment (native) | -0.60 | 0.58 | -1.04 | 44 | 0.296 |
| | Deer activity | -0.44 | 0.25 | -1.74 | 44 | 0.081 |
| | Trt (native)*Deer activity | 0.40 | 0.35 | 1.16 | 44 | 0.247 |
